# Analysis of cell-cell bridges in *Haloferax volcanii* using Electron cryo-tomography reveal a continuous cytoplasm and S-layer

**DOI:** 10.1101/2020.09.30.320622

**Authors:** Shamphavi Sivabalasarma, Hanna Wetzel, Phillip Nußbaum, Chris van der Does, Morgan Beeby, Sonja-Verena Albers

## Abstract

Halophilic archaea exchange DNA and proteins using a fusion-based mating mechanism. Scanning electron microscopy previously suggested that mating involves an intermediate state, where cells are connected by an intercellular bridge. To better understand this process, we used electron cryotomography and fluorescence microscopy to visualize cells forming these intercellular bridges. Electron cryo-tomography showed that the observed bridges were enveloped by an S-layer and connected mating cells via a continuous cytoplasm. Macromolecular complexes like ribosomes and unknown thin filamentous helical structures were visualized in the cytoplasm inside the bridges, demonstrating that these bridges can facilitate exchange of cellular components. We followed formation of a cell-cell bridge by fluorescence time-lapse microscopy between cells at a distance of 1.5 µm. These results shed light on the process of haloarchaeal mating and highlight further mechanistic questions.

## Introduction

Horizontal gene transfer is fundamental to archaeal and bacterial evolution. The diverse mechanisms of horizontal transfer, however, remain incompletely understood (Wagner, Whitaker, Krause, Heilers, Van Wolferen, et al. 2017). These mechanisms include uptake of DNA via natural transformation, transfer of conjugative plasmids, transduction, uptake of DNA via membrane vesicles and cell fusion hybrids (Wagner, Whitaker, Krause, Heilers, van Wolferen, et al. 2017). Members of Euryarchaeota, *Pyrococcus furiosus* and *Thermococcus kodakaraensis* are naturally competent taking up linear and circular DNA (Lipscomb et al. 2011; Sato et al. 2005). Transfer of conjugative plasmids was described first in Sulfolobales by the isolation of the first archaeal conjugative plasmid in 1995 (Prangishvili et al. 1998; Schleper et al. 1995; Stedman et al. 2000). Interestingly, analysis of the genome of Sulfolobales revealed the insertion of proviral DNA from Sulfolobus spindle-shaped virus 1 (SSV1) (Schleper, Kubo, and Zillig 1992). SSV1 stays integrated in archaeal genomes and producing viral particles budding from the cells for the transfer of viral DNA (Quemin et al. 2016). *Methanococcus voltae* PS produces viral particles named “voltae transfer agent” (VTA) which can carry chromosomal fragments instead of viral DNA (Bertani 1999; Eiserling et al. 1999; Lang, Zhaxybayeva, and Beatty 2012). Similarly to VTA, *Thermococcales* release membrane vesicles packed with chromosomal and plasmid DNA for the exchange of genetic material (Soler et al. 2008). Members of *Sulfolobus spp*. can exchange DNA upon UV-induced DNA damage allowing for DNA repair using homologous recombination (Ajon et al. 2011; Fröls et al. 2008). Cell aggregates are formed mediated by UV-induced pili (Ups-pili) and the crenarchaeal exchange of DNA system (Ced-system) is activated (Ajon et al. 2011; Fröls et al. 2008; Fröls, White, and Schleper 2009). Using the Ups-pili cell-cell contact is established and DNA is exchanged (Van Wolferen et al. 2016). Remarkably the exchange is species-specific possibly being mediated by the degree of N-glycosylation of Ups-pili (van Wolferen et al. 2020). Finally bidirectional gene transfer occurs in haloarchaea via cell fusion (Mevarech and Werczberger 1985; Rosenshine, Tchelet, and Mevarech 1989).

Here, the bidirectional DNA transfer through cell fusion in haloarchaea is further elucidated. In the 1980s, it was described that mixing of two different auxotrophic strains of the halophilic euryarchaeon *Haloferax volcanii*, resulted in prototrophic recombinant cells. The mating frequency was determined in the presence of DNase to rule out natural transformation and was 10^−6^ (Mevarech and Werczberger 1985). It was proposed that transfer of genetic material occurred via an uncharacterized fusion-based mating mechanism (Mevarech and Werczberger 1985). Remarkably, the transfer of DNA in *H. volcanii* is bidirectional without a specific donor or recipient and since mating and subsequent DNA exchange has been observed between the geographically distant species *H. volcanii* and *H. mediterranei*, it is not necessarily species specific (Naor et al. 2012). It was observed that two *Haloferax volcanii* cells can fuse to form a hybrid state (Naor et al. 2012; Naor and Gophna 2013). In this state, large chromosomal DNA fragments are exchanged and after recombination followed by cell separation, this results in genetic hybrids of the parents (Naor et al. 2012; Naor and Gophna 2013). CRISPR spacers matching chromosomal genes, including housekeeping genes, are also exchanged between species (Turgeman-Grott et al. 2019). Strikingly, mating frequency depends on factors that impact the cell surface such as external salt concentration and N-glycosylation of the S-layer (Shalev et al. 2017). Defects in the N-glycan of S-layer proteins significantly reduce mating frequencies, suggesting an important role for S-layer glycosylation in initiation of cell-cell interaction and cell fusion (Shalev et al. 2017). Scanning electron micrographs of *H. volcanii* have suggested the formation of intermediate intercellular bridges prior to cell fusion (Rosenshine, Tchelet, and Mevarech 1989). These cell-cell bridges might allow for an exchange of genetic material and drive cell fusion (Naor and Gophna 2013). Exchange of genetic material has only been observed on solid media in previous studies, prompting questions about the mechanisms involved in mating. Formation of possible cell-cell bridges between cells has also been observed in other archaeal lineages, such as members of Sulfolobales (Schleper et al. 1995), Thermococcales (Kuwabara et al. 2005) and even between Nanoarchaea and Thermoplasmatales (Comolli and Banfield 2014). Formation of cell-cell bridges was also reported in bacterial species. These nanotubes are enveloped by a membrane layer and build a bridge between two neighboring bacterial cells allowing an exchange of cytoplasm (Baidya et al. 2018; Dubey and Ben-Yehuda 2011).

To better characterize the mechanism of horizontal gene transfer by fusion in *H. volcanii*, we used electron cryotomography (cryoET) to preserve whole cells in a frozen hydrated state. We identified and imaged cell-cell bridges connecting the cytoplasms of pairs of cells grown in liquid media. Tomograms revealed that two mating cells shared a continuous membrane, a continuous S-layer and had continuous connected cytosols. Strikingly, macromolecular structures were detected in the cell-cell bridges likely to be ribosomes. Fluorescence time-lapse microscopy of *H. volcanii* cells with fluorescently stained S-layers showed how cells established an intercellular bridge as an intermediate state prior to cell fusion.

## Results

### Whole cell *in situ* electron cryo-tomography captures cell-cell bridges between two *H. volcanii* cells

To investigate the structure of the cell-cell bridges in *H. volcanii*, whole-cell electron cryo-tomography (cryoET) was used. CryoET offers the possibility to image cells in their native environment in a near-native frozen-hydrated state to macromolecular resolutions. Cryomicrographs of vitrified *H. volcanii* cells grown in liquid medium were acquired. The initial tomograms showed archaellated cells with a possible storage granule as well as ribosomes in the cytoplasm (Figure 1). All detected cells were enveloped by a continuous paracrystalline and proteinaceous surface layer (S-layer) over the membrane (Figure 1). In *H. volcanii*, the S-layer consist of one highly glycosylated protein that is secreted and lipid anchored to the membrane (Kessel et al. 1988; Sumper et al. 1990). The S-layer protein self-assembles to a 2D-lattice around the cell acting as a molecular sieve, involved in cell recognition and cell shape maintenance (Sára and Sleytr 2000; Sleytr et al. 2014). Strikingly, S-layer proteins could be detected, arranged in a hexagonal lattice around the cell similarly as reported in an early study (Figure 1B) (Kessel et al. 1988). Upon closer investigation of a subtomogram slice, the dome shape morphology formed by S-layer proteins can be identified (Figure 1C) (Kessel et al. 1988). The thickness of the S-layer was determined by measuring the distance from the membrane to the S-layer protein. The average thickness was determined to 20.4 ± 2.7 nm (Supplementary Table S1).

**Figure 1:**
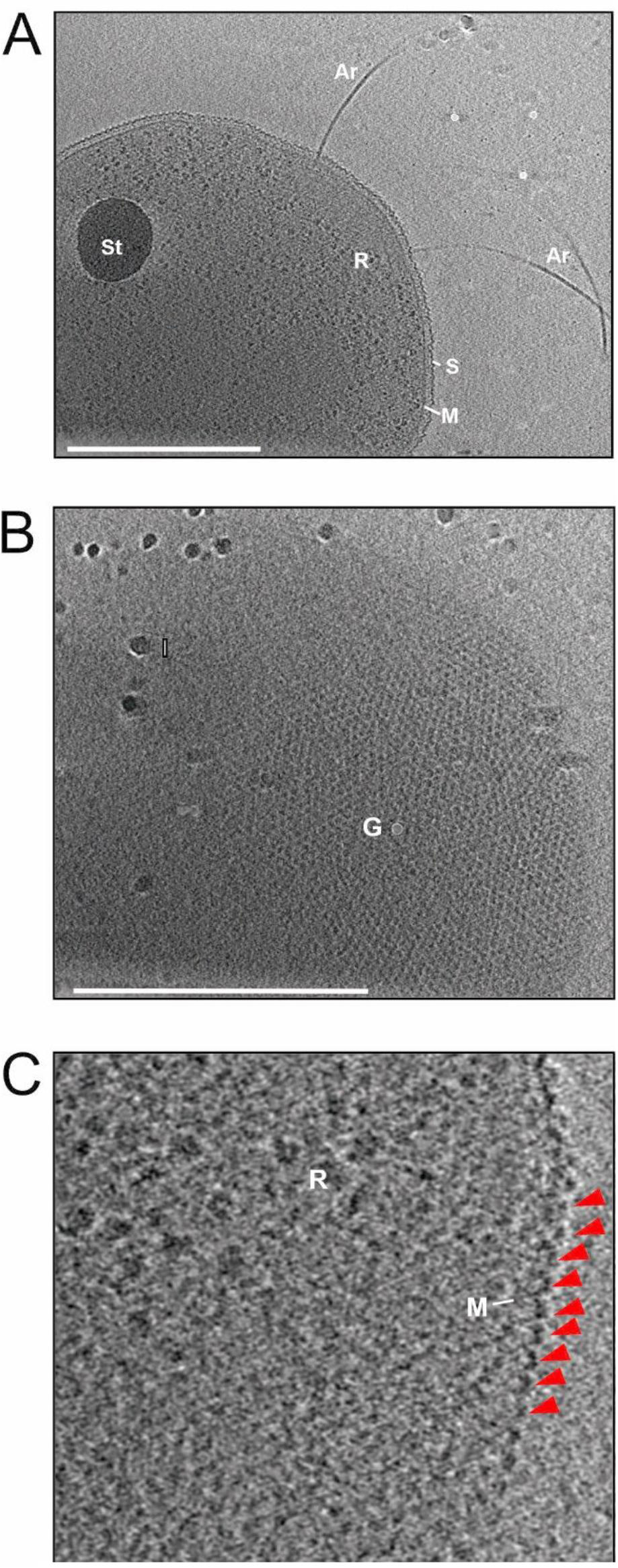
Electron cryo microscopy and subtomogram slices of *H. volcanii*. (A) A slice through the reconstructed tomogram of *H. volcanii*. The subtomographic slice reveals the cells filled with ribosomes (R) and a possible storage granule (St). The cells are enveloped by the cytoplasmic membrane (M) and a S-layer (S). From the cells, the archaellar motility system, the archaellum, extends (Ar). Scale bare is 500 nm. (B) Extracted subtomographic slice, showing the top view of the cell. The cell body is covered by the hexagonal arranged S-layer lattice. Gold fiducial (G) and ice crystal (I) contaminations are indicated. Scale bar is 500 nm. (C) A magnified tomographic slice from (A) showing the characteristic dome-like shape the arranged S-layer protein indicated by a red arrow. Scale bars in 100 nm.

Interestingly, cryoET allowed the observation of several cells in a hemifusion state connected via cell-cell bridges (Figure 2, upper panels). In total, out of 280 collected tilt series 20 tilt series of cell-cell bridges were acquired with a magnification sufficient to focus on the intercellular bridges (Figure 2, lower panels). As well as cell-cell bridges between intact cells (Figure 2, left panels), we observed cell bridges between intact and broken cells (Figure 2, middle panels) and disrupted cell bridges. These broken cells and disrupted bridges were probably ruptured during the blotting procedure. A representative tilt series is shown in supplementary movie S1.

**Figure 2:**
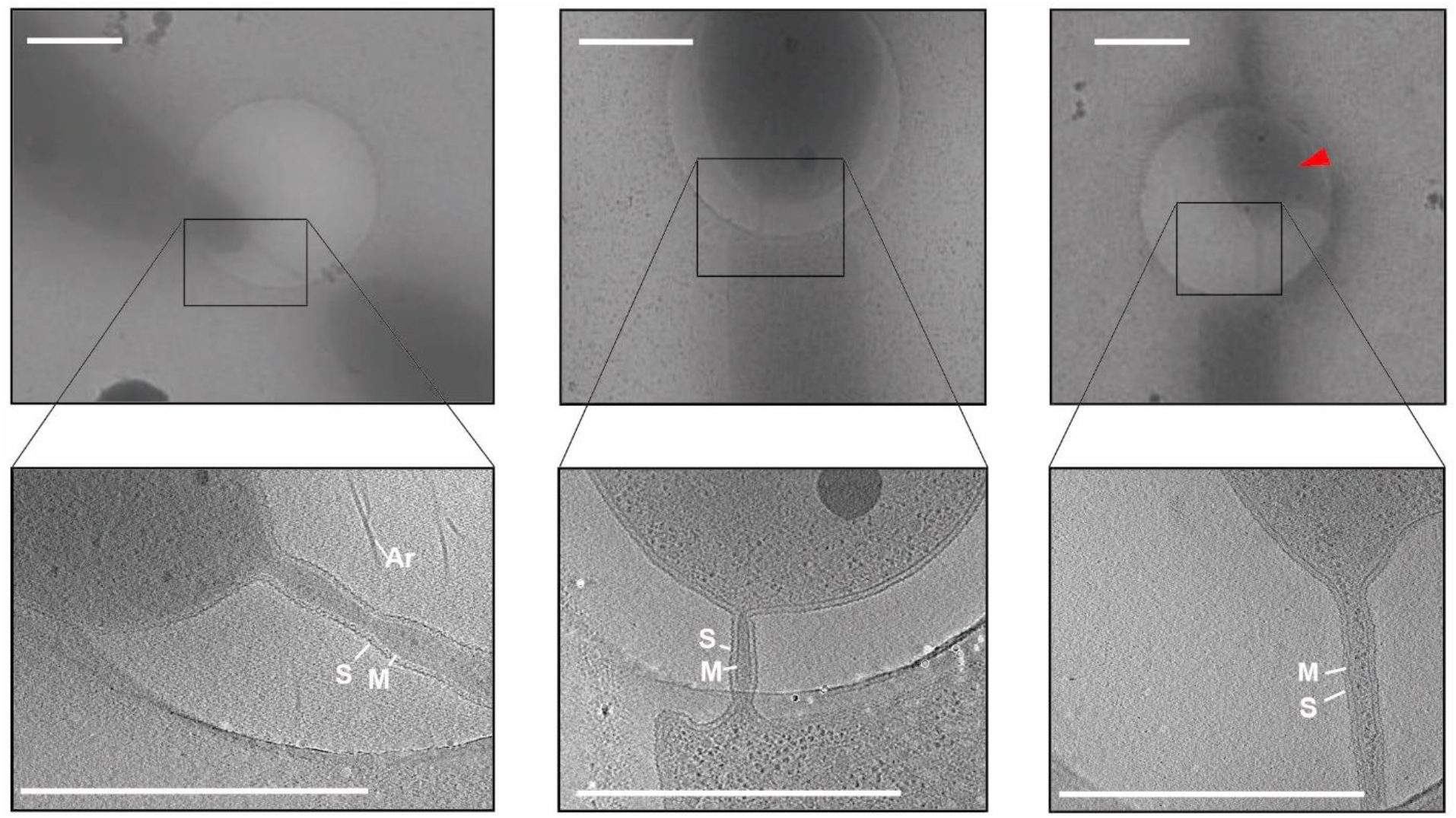
Electron cryo microscopy and tomographic slices of cell-cell bridges in *H. volcanii*. Low magnification cryo micrographs of cell-cell bridges targeted for cryoET (upper panels) with the corresponding tomographic slice from a reconstructed tilt series (lower panels). The electron cryo micrographs as well and the corresponding slice through tomograms depict the variety in shape, length and width of the cell-cell bridges. The right panel shows a partially ruptured cell indicated by a red arrow. Scale bars are 1 µm. The cytoplasmic membrane (M), the S-layer (S), and Archaella (Ar) are indicated.

We measured the widths and lengths of cell-cell bridges. The width was measured over the length of the cell-cell bridge and average width of each cell-cell bridge was determined (Figure 3, Table S1). For determination of the length, only the cell-cell bridges were considered that connected two cells as shown in Figure 2 (left and middle panel). The width varied from 57-162 nm and the length varied from 253-2144 nm (Figure 3, Table S1). The scatter plot and histogram (Figure 3) shows that most cell-cell bridges have a width up to 100 nm with a maximal length of 1 µm-1.2 µm indicating that cells might need to be within ∼1.2 µm for the formation of cell-cell bridges to occur. No relation between length and diameter could be detected.

**Figure 3:**
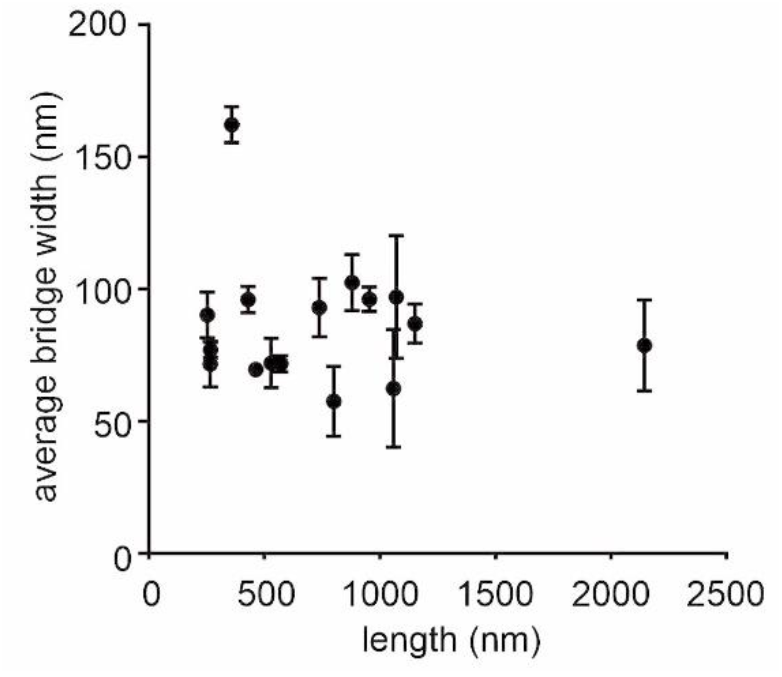
Determination of length and width of cell-cell bridges. (A) Scatter plot of length against the corresponding average width of cell-cell bridges. Only cell-cell bridges were considered that connected two cells together.

### Cell-cell bridges are surrounded by a continuous S-layer and connect the cytoplasms of two cells

Closer investigation of the cell-cell bridges showed that these bridges connected the cytoplasms of the two cells. The connected cytoplasm was surrounded by a continuous cytoplasmic membrane and a continuous S-layer. (Figure 4). Since the cytoplasms of the two cells are connected, this would allow exchange of cytoplasmic materials between two cells. Indeed, we saw large macromolecular complexes consistent with the size, shape, and density of ribosomes within the tubular cell-cell bridges (Figure 4, Movie S1), suggesting that high molecular weight complexes are exchanged between mating cells. Next to the ribosomes also unknown thin filamentous helical structures of 199 ± 18 nm long and 9.0 ± 2.3 nm wide (Figure 4B, Movie S1) were observed in the one of the analyzed cell-cell bridge. The function and the proteins that form the thin filamentous helical structures are unclear although we speculate they may be cytomotive filaments to drive cytoplasmic exchanges.

**Figure 4:**
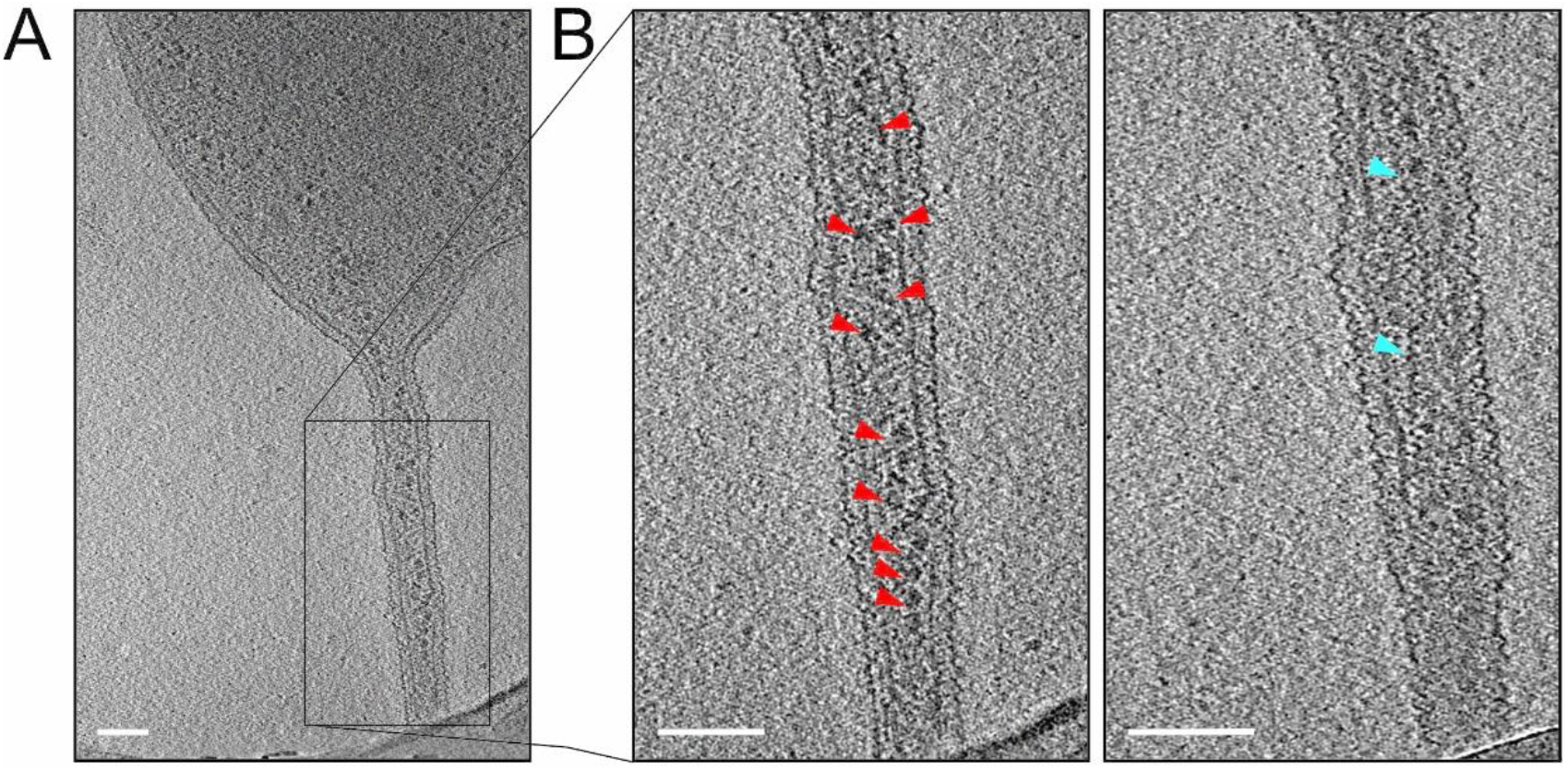
Detection of macromolecular complexes in cell-cell bridges. (A) A micrograph of a targeted cell-cell bridge shows the presence of ribosomes in the cell-cell bridge (as indicated by red arrows). (B) The left panel shows the magnified tomographic slice of the cell-cell bridge and shows the ribosomes arranged in a chain-like manner (as indicated by red arrows). The right panel shows another slice of the selected cell-cell bridge showing a long filamentous structure inside the cell-cell bridge (as indicated by blue arrows). Scale bars are 100 nm

### *In vivo* observation of the formation of an intercellular *H. volcanii* cell-cell bridge

The tomograms showed that intercellular bridges are encapsulated by an S-layer. To follow the formation of a cell-cell bridge using time-lapse fluorescence microscopy, the cells were incubated with the Alexa Fluor 488 NHS Ester and the cells were followed over a period of 16 hours. As expected, the cells were mainly fluorescently labelled on the outside. Comparison of the proteins which were labelled with the fluorescent probe in the total cell extract with isolated S-layers showed that, next to the S-layer protein, also several other proteins were labelled (Supplementary figure 1). Several experiments were conducted where over 16 hours every 30 min fluorescent and phase contrast images were acquired. In one of these experiments, the formation of a cell-bridge was observed. Notably, after five hours of incubation, one thin fluorescent connection between two cells was detected that can be identified as a *de novo* formed cell-cell bridge. (Figure 5, Movie S2). The time lapse movie shows fluorescent cells with increasing cell size due to an unknown cell division defect where a fluorescent septum is formed between two adjacent cells which do not separate. This is sometimes observed during these experiments and is most likely unrelated to the observed cell-cell bridge. Between the 2:30 and 3:00 hr time points a full cell-cell bridge is formed. Remarkably, the cell-cell bridge is formed between two cells at 1.5 µm distance without any initial direct contact, suggesting that cell-cell bridge formation is an active process. CryoET showed that shortly after cell-cell bridge formation the cytoplasms are most likely connected and transfer between the two cells can occur through the cell-cell bridges. It was not possible to determine whether the cell-cell bridge is formed from one or from both cells. After 7.5 hours, the length of the cell-cell bridge is decreasing indicating a contraction to bring both cells in close proximity (Figure 5). The cell-cell bridge was maximally 1.5 µm long, growing shorter and thicker over time. Unfortunately, the *in vivo* formation of only one cell-cell bridge could be observed as cell-cell bridge formation, however, this is the first time that the formation of a cell-cell bridge could be observed in real time.

**Figure 5:**
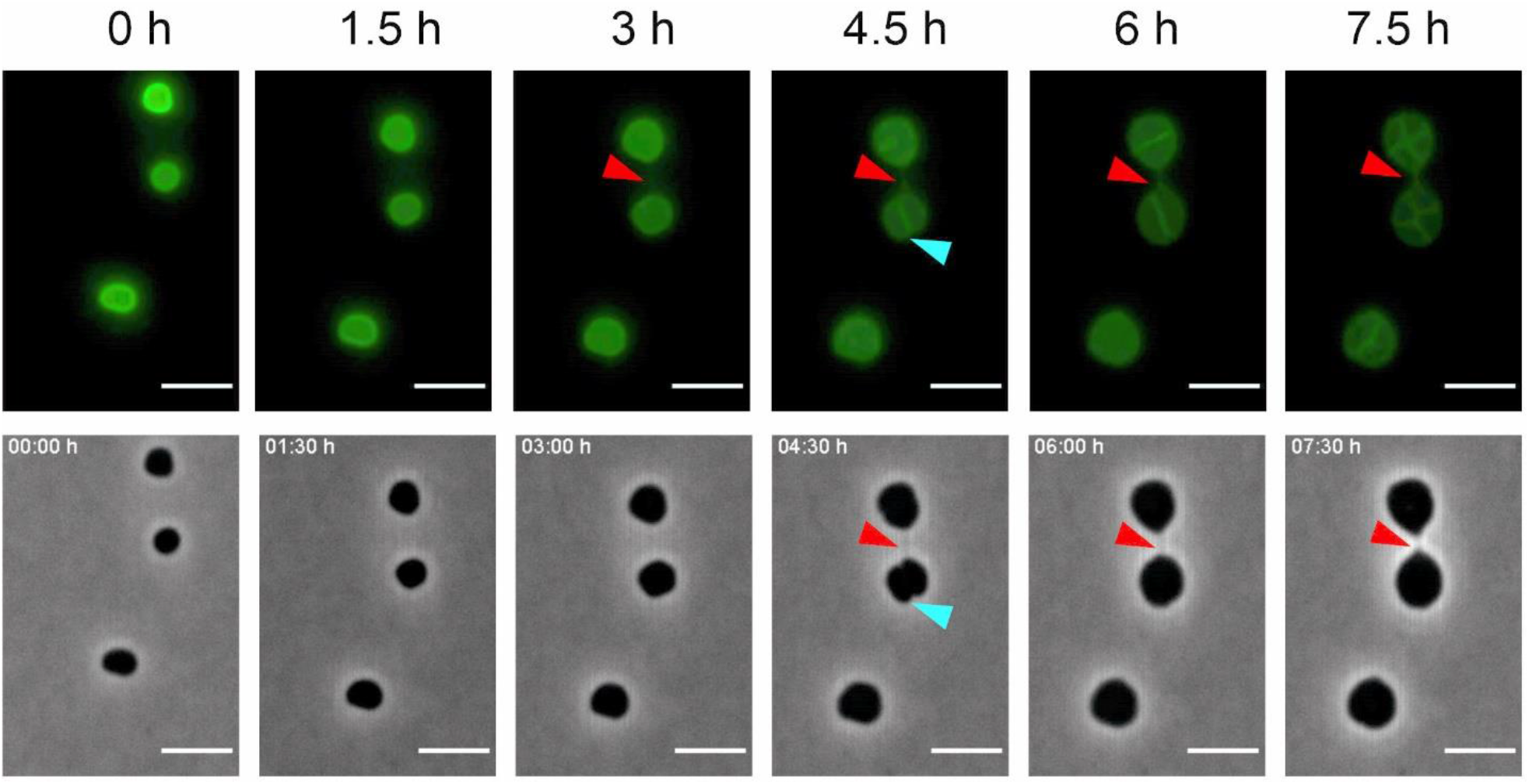
Formation of a cell-cell bridge followed by fluorescence microscopy. Time-lapse fluorescence images of AlexaFluor488 labeled *H. volcanii* cells with the corresponding phase-contrast image. The fluorescence signal for the cell-cell bridge can be detected after 5 h and is indicated by a red arrow. In phase contrast, the cell-cell bridge can be detected after 7 h. Blue arrows indicate the septum formed between the cells due to a cell division defect. Scale bars are 4 µm.

## Discussion

Mechanisms of gene transfer are diverse in Archaea (Ajon et al. 2011; Allers 2011; Bertani 1999; Meile, Abendschein, and Leisinger 1990; Naor et al. 2012; Prangishvili et al. 1998; Schleper et al. 1995; van Wolferen et al. 2013, 2020). In *H. volcanii* genetic transfer occurs in a bidirectional manner upon fusion (Mevarech and Werczberger 1985; Naor and Gophna 2013; Rosenshine, Tchelet, and Mevarech 1989) and it was proposed that the formation of cell-cell bridges may precede fusion as an intermediate state (Naor and Gophna 2013; Rosenshine, Tchelet, and Mevarech 1989).

Whole-cell cryoET enabled preservation and the study of cells in this intermediate bridged state, while time-lapse fluorescence allowed for the observation of the formation of cell-cell bridges in real-time. Despite the high salinity of the cell cytoplasm and the media, electron cryo-tomograms gave a high resolution snapshot of the features of the cell-cell bridges in *H. volcanii*. However, only a limited number of cell-cell bridges could be studied. In contrast to the cell-cell bridges observed by Mevarech and Rosenshine on solid media, bridge formation and fusion events were recorded in liquid media, but in liquid media, these events occur most likely much less often than on solid media (Rosenshine, Tchelet, and Mevarech 1989). Furthermore, more cell-cell bridges would probably have been observed in an medium containing a higher salt concentration, as mating efficiency was shown to be higher at 3.4 M NaCl instead of the 2.2 M NaCl in the medium used (Shalev et al. 2017). However higher salt would reduce the clarity of electron micrographs.

Time-lapse fluorescence microscopy allowed for the observation of the *de novo* formation of a cell-cell bridge. This showed that cells are able to bridge the distance by the formation of a cell-cell bridge reaching to the adherent cell. The time-lapse fluorescence microscopy showed a cell-cell bridge that was formed over a distance of 1.5 µm whereas the cryoET showed that most cell-bridges were shorter than 1-1.2 µm, which demonstrates that cell-bridges with a length of ∼1 µm can be easily formed. Full cell-cell bridge formation was observed within 0.5 hr, suggesting it is a relatively fast process (See movie S2 between 2:30 and 3:00 hrs). Indeed, a recent transcriptomic study showed that mating impacts genes involved in cell division and glycosylation. Strikingly, increased expression was detected for selfish genetic elements, restriction-modification system and CRISPR-Cas. These changes were detected in the first hours after transfer to a filter paper, indicating that mating efficiency is probably the highest 4 to 8 h after cell contact (Makkay et al. 2020). Since adherence must be the first step before fusion, cells may initially form an S-layer covered protrusion that detects the adherent cell and initiates formation of the cell-cell bridge. Indeed, glycosylation of the S-layer is crucial in intraspecies mating efficiency (Shalev et al. 2017). Significantly fewer fusion events were observed when neither tetra-nor pentasaccharides decorated the S-layer protein (Shalev et al. 2017). In *Sulfolobales*, N-glycosylation of UV-inducible pili is also crucial for species-specific UV-induced aggregation. Addition of different sugars to the media leads to decreased aggregation formation and ensuing reduced DNA exchange for DNA repair (van Wolferen et al. 2020). Similar to *Sulfolobales*, the N-glycan of a cell might be detected by surface-expressed receptor proteins for cell-cell recognition prior to cell-cell bridge formation.

All cell-cell bridges that were detected contained a connected S-layer and cytoplasm suggesting that the fusion of the cytoplasms occurs very shortly after the cell-cell bridge is formed. Time-lapse fluorescence microscopy indicated that the cell-cell bridge may shorten and widen for subsequent fusion of two cells (Figure 5, Movie S2). The electron cryo-tomograms revealed a variety in the size of cell-cell bridges possibly due to the different fusion states that were detected. Upon closer investigation of the cell-cell bridges, ribosomes and other complexes among which thin filamentous helical structures were detected indicating an exchange of cytoplasmic components.

Cell-cell bridges (nanotubes) have also been identified between members of bacterial species and also connect the cytoplasms of neighboring cells, but still little is known about how they are formed (Baidya et al. 2018; Dubey and Ben-Yehuda 2011). In Eukarya, cell fusion is a common mechanism, where gametes, myoblasts, or vesicles fuse to a partner or host cell. Fusion events are mostly mediated via fusogens (SNARE proteins) or fusexins in eukaryotic cells (Hernández and Podbilewicz 2017; Segev, Avinoam, and Podbilewicz 2018). By a controlled re-organization/folding of the fusogens or fusexins in the membranes of the mating cells, the high energy barrier to cell-cell fusion can be overcome (Segev, Avinoam, and Podbilewicz 2018). Although 2 DedA-like proteins related to the SNARE-associated Tvp38 proteins in eukaryotes have been identified in *H. volcanii*, their role in the formation of cell-cell bridges remains unclear. Also the role and identity of the thin filamentous helical structures observed in the cell-cell-bridges remains unknown.

Here we have shown that *H. volcanii* can form cell-cell bridges which contain a continuous cytoplasm through which large molecular complexes like ribosomes can be exchanged. The formation of a cell-cell bridge was observed between two cells ∼1.5 um apart, suggesting this is an active process. These observations raise several new questions which should be addressed in future studies; How are these cell-cell bridges formed, do these cell-cell bridges grow from one or from both cells, how do the cell-cell bridges fuse?, and which proteins are required for the formation of the cell-cell bridges ?

## Material and Methods

### Strains and growth conditions

Growth of *H. volcanii* H26 and RE25 was performed as described previously (Allers et al. 2004; Esquivel and Pohlschroder 2014). The cells were grown in YPC medium with yeast extract, peptone and casamino acids (Bacto) or selective Casamino acid medium (CA medium) and CA medium supplemented with trace elements for selective growth as described (CAB) (Duggin et al. 2015).

### Electron cryo-tomography

*H. volcanii RE 25* was inoculated in 5 mL CA media plated supplemented with 1 g/l thiamine and 0.1 µg/L biotin, 10 µg/mL uracil and 50 µg/mL tryptophan that was incubated at 42 °C overnight. 5 µL and 15 µL of the pre-culture was inoculated in 20 mL CA media and incubated again at 42 °C overnight. At OD_600_ of 0.05, the cells were harvested at 2000 x g for 20 min at 40 °C. The pellet was dissolved in 1 mL CA media and again pelleted at 2000 x g for 10 min at 40 °C. The cell pellet was dissolved again in CA medium to a theoretical OD_600_ of 3 or 5. The cells were mixed with BSA-coated 10 nm gold fiducial markers and 2.5 µL of cells were applied to a freshly glow-discharged copper Quantifoil R2/2 grid (300 mesh). The vitrification of the grid was done using the Vitrobot Mark IV (FEI). The grid was blotted on the back using a repellent Teflon membrane and subsequently plunge-frozen in liquid ethane. Electron cryotomography was conducted with a FEI F20 (FEG) equipped with a Falcon II direct electron detector. A total cumulative electron dose of 120 e-/Å^2^ was used per tilt series with −3 to −6 µm defocus using a tilt range of ± 53 ° with 3 ° increments and a pixel size of 8.28 Å. Data was acquired using Leginon (Carragher et al. 2000). Tomograms were reconstructed automatically using RAPTOR software and IMOD (Amat et al. 2008; Kremer, Mastronarde, and McIntosh 1996; Mastronarde 1997).

### S-layer staining

*H. volcanii* H26 was inoculated in 5 mL CAB medium supplemented with 10 µg/mL uracil and grown overnight at 45 °C. 220 µL were inoculated in 20 mL of the main culture which was grown overnight at 45 °C. The cells were harvested at OD_600_=0.2 at 1800xg for 10 min at 25 °C. The pellet was resuspended in 2 mL of buffered media and washed three times at 3400xg for 10 min at RT and resuspended in 500 µL buffered media. The pH was adjusted to pH 8-8.5 with 1 M NaHCO_3_ and 50 µg of Alexa Fluor 488 NHS Ester (Thermo Fisher Scientific) was added. The cells were incubated at RT for 1h while rotating. To remove excess dye, the cells were washed three times as above with 500 µL CAB media.

### Isolation of stained S-layer

Staining was checked by sonicating the cells for 10 min in an ultrasonic bath. Afterwards, the cell debris was pelleted at 3400xg for 10 min at RT. SDS was added to a final concentration of 0.01 % and the mixture was centrifuged again at 3400xg for 10 min at RT. The supernatant was divided into 2x 500 µL and centrifuged at 190000xg for 1h at 4 °C. The resulting pellets were resuspended in 50 µl 1x loading dye and in 5 µL 1x PBS, 0.1 % Triton-X 100. Both pellets were incubated at 6 °C, in a light-protected manner for 48h and then mixed together. This was incubated for 2 1/2 h at 37 °C and 10 µL was used for an SDS-PAGE and detection of the fluorescence signal.

### S-layer isolation

For S-layer isolation, 400 mL of H26 was grown at 45 °C to an OD_600_ of 1.37. The cells were pelleted at 6200xg for 25 min at 4 °C. The pellet was resuspended in 200 mL CA media and 60 mL of 0.5 M EDTA (pH 6.7) was added. Subsequently, the pellet was incubated at 37 °C while shaking for 30 min. The spheroplasts were removed via centrifugation in an iterative manner at 3000xg for 15 min, 7000xg for 5min and 13 000xg for 10 min. The supernatant was concentrated via Amicon (MWCO= 50kDa, Merck Millipore) to 500 µL. 16 µL were used for an SDS-PAGE and sent for Mass spectrometry.

### Fluorescence time-lapse microscopy

For microscopy, 3 µL of stained cells were pipetted on a 0.3 % agarose pad consisting of agarose dissolved in CAB medium supplemented with 10 µg/mL uracil. Phase-contrast and fluorescence images were captured every 30 minutes for 16h at 100x magnification at pH 3 and GFP mode using a Zeiss Axio Observer 2.1 microscope equipped with a heated XL-5 2000 Incubator while running VisiVIEW^®^ software. Images were analyzed using ImageJ, Fiji (Schindelin et al. 2012).

## Data Availability Statement

The raw data will be made available by the authors, without undue reservation.

## Author contributions

SS, PN, MB and SVA designed research. SS performed cryoET with technical help from FR. SS, SVA and MB analysed the tomograms and designed figures. HW and PN performed fluorescence microscopy. HW, PN, SS and SVA analysed the fluorescence microscopy data and designed figures. SS, MB and SVA wrote the manuscript. All authors read and reviewed the manuscript.

## Acknowledgments

We thank Dr Florian Rossmann and Paul Simpson for technical assistance with imaging *H. volcanii* for CryoET in the Imperial College Centre for Structural Biology Electron Microscopy Facility.

