## Supplementary Material for "Analysis of cell-cell bridges in *Haloferax volcanii* using Electron cryo-tomography reveal a continuous cytoplasm and S-layer"

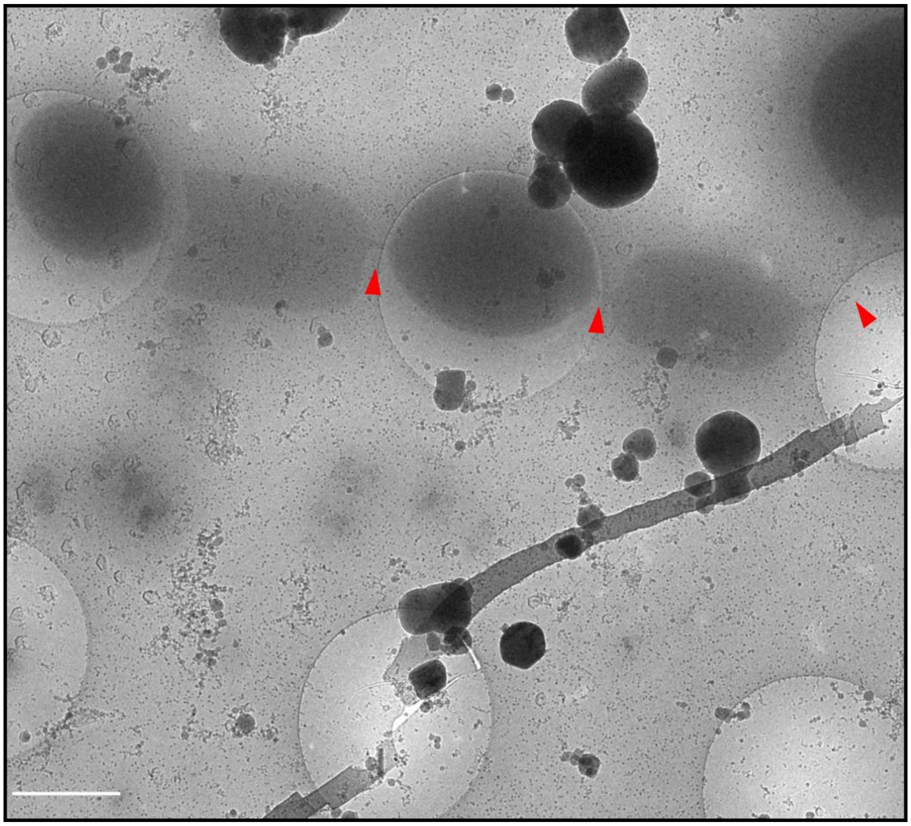

**Supplementary Figure 1: Cryo-electron micrograph of a chain of cells.** The cell body are connected via intercellular bridges indicated with the red arrows. Scale bar is 500 nm.

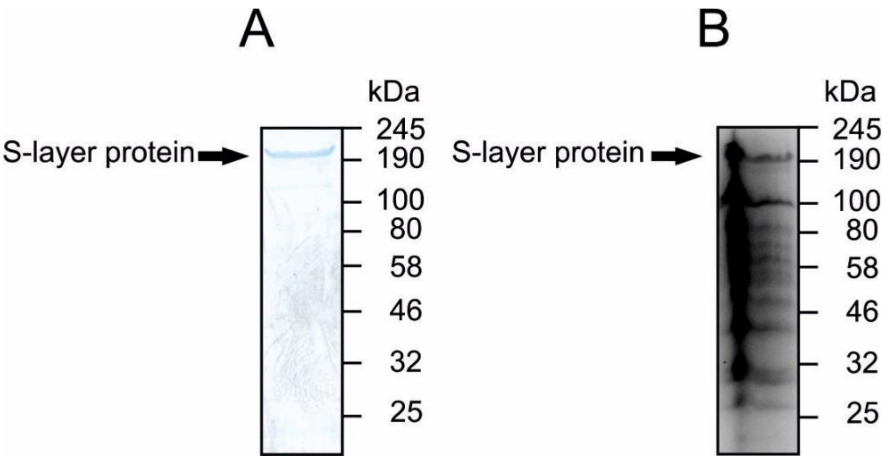

**Supplementary Figure 2: SDS-PAGE and Fluorescence detection of isolated S-layer from *H. volcanii.*** (A) To verify the staining of the S-layer with AlexaFluor488, S-layer was isolated. The indicated band running at 190 kDa was verified as S-layer glycoprotein via Mass spectrometry. (B) The S-layer and membrane were isolated from fluorescently stained cells and the fluorescence signal was detected. The band running at 190 kDa corresponds to (A) and can be determined as S-layer glycoprotein.

**Supplementary movie 1:**

**Tomographic reconstruction of a cell-cell bridge. Scale bar is 100nm**

**Supplementary movie 2:**

**Fluorescence time-lapse microscopy of *H.volcanii* cells on an agar pad stained with Alexa Fluor 488 NHS Ester. Scale bar is 4 µm**

**Supplementary Table 1: Measured S-layer thickness and cell-cell width of selected reconstructed tomograms**

| Session and tomography stack number | | S-Layer /nm | Average thickness S-Layer /nm | Bridge width/nm | | Average bridge of width /nm | Standard error width | Length /nm |
|---|---|---|---|---|---|---|---|---|
| 1 | 1 | 16.80 | 15.20 | 107.39 | 81.23 | 96.65 | 23.24 | 1070.96 |
| | | 14.64 | | 107.75 | 61.47 | | | |
| | | 14.16 | | 127.96 | 94.09 | | | |
| | 7 | 23.60 | 23.24 | 106.93 | 94.38 | 93.32 | 11.06 | 738.71 |
| | | 22.90 | | 96.14 | 74.63 | | | |
| | | 23.02 | | 99.75 | 88.11 | | | |
| | | 23.42 | | | | | | |
| | 20 | 18.73 | 20.46 | 79.63 | 75.56 | 77.10 | 2.96 | 266.94 |
| | | 20.04 | | 80.05 | 72.78 | | | |
| | | 22.61 | | 79.24 | 75.34 | | | |
| | 30 | 14.48 | 17.58 | 100.19 | 97.12 | 96.43 | 5.05 | 429.33 |
| | | 20.70 | | 96.93 | 102.73 | | | |
| | | 17.55 | | 88.63 | 92.97 | | | |
| | 37 | 22.60 | 19.25 | 82.21 | 85.96 | 86.54 | 7.43 | 1151.52 |
| | | 16.30 | | 82.10 | 81.49 | | | |
| | | 18.84 | | 101.10 | 86.40 | | | |
| 2 | 15 | 21.73 | 19.60 | 97.43 | 99.93 | 102.44 | 10.62 | 880.48 |

| | | | | | | | | |
|---|---|---|---|---|---|---|---|---|
| | | 17.83 | | 117.01 | 118.44 | | | |
| | | 19.25 | | 92.87 | 108.40 | | | |
| | 23 | 17.19 | 19.26 | 90.61 | 65.33 | 78.75 | 17.31 | 2144.91 |
| | | 20.69 | | 103.77 | 61.26 | | | |
| | | 19.91 | | 86.44 | 65.11 | | | |
| | 29 | 24.61 | 22.63 | 79.48 | 76.33 | 71.67 | 8.68 | 265.32 |
| | | 21.49 | | 61.32 | 81.21 | | | |
| | | 21.78 | | 62.11 | 69.57 | | | |
| | 41 | 23.33 | 22.30 | 57.96 | 80.38 | 72.00 | 9.43 | 529.35 |
| | | 21.73 | | 72.27 | 72.02 | | | |
| | | 21.85 | | 85.77 | 72.43 | | | |
| | 56 | 27.12 | 25.71 | 81.70 | 49.05 | 57.63 | 13.33 | 801.29 |
| | | 26.00 | | 54.06 | 43.05 | | | |
| | | 24.01 | | 57.10 | 60.80 | | | |
| | 73 | 19.54 | 19.81 | 168.34 | 169.09 | 162.24 | 6.81 | 358.65 |
| | | 20.24 | | 163.55 | 152.62 | | | |
| | | 19.66 | | 154.83 | 161.20 | | | |
| | 87 | 18.51 | 18.32 | 98.43 | 98.72 | 96.12 | 4.69 | 955.13 |
| | | 17.12 | | 99.21 | 88.94 | | | |
| | | 19.32 | | 90.72 | 91.97 | | | |
| 3 | | | | | | | | |

| | | | | | | | | |
|---|---|---|---|---|---|---|---|---|
| | 67 | 18.86 | 23.18 | 148.05 | 124.85 | 131.46 | 10.54 | - |
| | | 26.24 | | 121.20 | 131.98 | | | |
| | | 24.43 | | 123.22 | 139.44 | | | |
| | 53 | 18.38 | 19.41 | 81.80 | 91.83 | 90.25 | 8.65 | 253.13 |
| 4 | | 19.60 | | 88.87 | 105.21 | | | |
| | | 20.24 | | 81.80 | 92.02 | | | |
| | 79 | 20.56 | 20.00 | 108.11 | 75.46 | 100.69 | 52.54 | - |
| | | 21.85 | | 113.27 | 132.93 | | | |
| | | 17.60 | | 163.16 | 11.20 | | | |
| | 146 | 19.78 | 20.40 | 71.23 | 67.33 | 71.77 | 3.04 | 570.92 |
| | | 20.30 | | 73.11 | 69.65 | | | |
| | | 21.13 | | 75.98 | 73.30 | | | |
| 5 | | | | | | | | |
| | 6 | 22.80 | 20.66 | 154.64 | 125.70 | 133.71 | 29.82 | - |
| | | 19.91 | | 133.51 | 105.63 | | | |
| | | 19.27 | | 180.34 | 102.41 | | | |
| | 61 | 16.41 | 19.05 | 149.66 | 162.43 | 152.09 | 43.24 | - |
| | | 19.53 | | 199.80 | 100.85 | | | |
| | | 21.21 | | 196.40 | 103.38 | | | |
| | 72 | 19.87 | 20.52 | 69.55 | 62.40 | 69.60 | 1.17 | 463.76 |

| | | | | | | | | |
|---|---|---|---|---|---|---|---|---|
| | | 21.53 | | 70.80 | 68.50 | | | |
| | | 20.15 | | 68.46 | 69.56 | | | |
| | 86 | 19.94 | 19.68 | 50.99 | 102.03 | 62.42 | 22.26 | 1059.26 |
| | | 20.21 | | 50.43 | 46.82 | | | |
| | | 18.88 | | 48.13 | 76.09 | | | |

**Supplementary Table 2: Measures average thickness of S-layer, widths of cell-cell bridges and measured lengths of the cell-cell bridges**

| Average S-layer thickness/ nm | average bridge width/nm | length/nm |
|---|---|---|
| 15.20 | 96.65 | 1070.96 |
| 23.24 | 93.32 | 738.71 |
| 20.46 | 77.10 | 266.94 |
| 17.58 | 96.43 | 429.33 |
| 19.25 | 86.54 | 1151.52 |
| 19.60 | 102.44 | 880.48 |
| 19.26 | 78.75 | 2144.91 |
| 22.63 | 71.67 | 265.32 |
| 22.30 | 72.00 | 529.35 |
| 25.71 | 57.63 | 801.29 |
| 19.81 | 162.24 | 358.65 |
| 18.32 | 96.12 | 955.13 |
| 23.18 | 131.46 | - |
| 19.41 | 90.25 | 253.13 |
| 20.00 | 100.69 | - |
| 20.40 | 71.77 | 570.92 |
| 20.66 | 133.71 | - |
| 19.05 | 152.09 | - |

| | | |
|---|---|---|
| 20.52 | 69.60 | 463.76 |
| 19.68 | 62.42 | 1059.26 |
